# Coupling of mouse olfactory bulb projection neurons to fluctuating odour pulses

**DOI:** 10.1101/2020.11.29.402610

**Authors:** Debanjan Dasgupta, Tom P.A. Warner, Andrew Erskine, Andreas T. Schaefer

**Affiliations:** The Francis Crick Institute, Neurophysiology of Behaviour Laboratory, London, UK; Department of Neuroscience, Physiology & Pharmacology, University College London, UK

## Abstract

Odours are transported by turbulent air currents, creating complex temporal fluctuations in odour concentration. Recently, we have shown that mice can discriminate odour stimuli based on their temporal structure, indicating that information present in the temporal structure of odour plumes may be extracted by the mouse olfactory system. Here using *in vivo* electrophysiological recordings, we show that mitral and tufted cells (M/TCs), the projection neurons of the mouse olfactory bulb, can encode the dominant temporal frequencies present in odour stimuli up to frequencies of at least 20 Hz. We show that M/TCs couple their membrane potential to odour concentration fluctuations; coupling was variable between M/TCs but independent of the odour presented and with TCs displaying slightly elevated coupling compared to MCs in particular for higher frequency stimulation (20Hz). Pharmacologically blocking the inhibitory circuitry strongly modulated frequency coupling. Together this suggests that both cellular and circuit properties contribute to the encoding of temporal odour features in the mouse olfactory bulb.

## Introduction

Temporal structure has long been considered as an integral part of sensory stimuli, notably in vision (Borghuis, Tadin, Lankheet, Lappin, & van de Grind, 2019; Buracas, Zador, DeWeese, & Albright, 1998; Chou, Toft-Nielsen, & Porciatti, 2019; Kauffmann, Bourgin, Guyader, & Peyrin, 2015; Kauffmann, Ramanoel, & Peyrin, 2014; Wang, Garcia, Sabottke, Spencer, & Sejnowski, 2019) and audition (Deneux, Kempf, Daret, Ponsot, & Bathellier, 2016; Nelken, Rotman, & Bar Yosef, 1999; Theunissen & Elie, 2014; VanRullen, Zoefel, & Ilhan, 2014). Odours in natural environments are transported by turbulent air streams, resulting in complex spatiotemporal odour distributions and rapid concentration fluctuations (Celani, Villermaux, & Vergassola, 2014; Pannunzi & Nowotny, 2019; Shraiman & Siggia, 2000). The neuronal circuitry of the olfactory system, particularly in invertebrates, has been shown to encode temporal structures present in odour stimuli (Hendrichs, Katsoyannos, Wornoayporn, & Hendrichs, 1994; Huston, Stopfer, Cassenaer, Aldworth, & Laurent, 2015; Pannunzi & Nowotny, 2019; Szyszka, Gerkin, Galizia, & Smith, 2014; Vickers & Baker, 1994; Vickers, Christensen, Baker, & Hildebrand, 2001). Temporal features in odour stimuli such as differences in stimulus onset were shown to be detectable on a behavioural level by bees (Sehdev, Mohammed, Triphan, & Szyszka, 2019; Szyszka, Stierle, Biergans, & Galizia, 2012). While the mammalian olfactory system is often considered a “slow” sense with single sniffs being the units of information (Cury & Uchida, 2010; Kepecs, Uchida, & Mainen, 2006; Wachowiak, 2011), recent reports have indicated that the neural circuitry of the early olfactory system readily sustains temporally precise action potential discharge and is capable to relay information about optogenetic stimuli with ~10 millisecond precision (Rebello et al., 2014; Shusterman, Smear, Koulakov, & Rinberg, 2011; Smear, Shusterman, O’Connor, Bozza, & Rinberg, 2011). Furthermore, we have recently shown that mice can utilise information in the temporal structure of odour stimuli at frequencies as high as 40 Hz to guide behavioural decisions (Erskine, Ackels, Dasgupta, Fukunaga, & Schaefer, 2019). However, how specific temporal features are represented in individual neurons in the brain remains unclear.

The olfactory bulb (OB) is the first stage of olfactory processing in the mammalian brain. Olfactory sensory neurons (OSNs) in the nasal epithelium convert chemical signals into electrical activity forming the input to the OB. They are known to show slow response profiles to odour stimuli (Sicard, 1986). This, together with the slow GPCR based odour-receptor interaction (Reisert & Matthews, 2001), created the general notion that mammalian olfaction has a limited temporal bandwidth. While OSN activity reflects only a low pass filtered version of the incoming odour signal, information about different frequency components might still be present in OSN activity (Nagel & Wilson, 2011). Moreover, circuit mechanisms in other brain regions and species have been shown to boost high-frequency content and sharpen stimulus presentation (Atallah & Scanziani, 2009; Nagel, Hong, & Wilson, 2015; O’Sullivan, Weible, & Wehr, 2019; Tramo, Shah, & Braida, 2002). Given the intricate circuitry present in the OB, where multiple types of interneurons (Burton, 2017) process incoming signals, we decided to investigate the role of the OB circuitry in representing and processing the temporal features of odour stimuli.

In this study, we show that mitral and tufted (M/T) cells, the OB output neurons, respond to odours - temporally modulated at frequencies of 2-20 Hz - in a frequency-dependent manner. Using whole-cell recordings, we show that subthreshold M/T cell activity *in vivo* can follow odour frequencies both at sniff and supra-sniff range for monomolecular odours and odour mixtures. We observe that while putative tufted (pTC) and mitral cells (pMC) show similar frequency coupling capacity at 2Hz, tufted cells have a higher propensity to follow odour frequencies at 20Hz. Pharmacologically “clamping” GABA receptors (Fukunaga, Berning, Kollo, Schmaltz, & Schaefer, 2012) we show that local inhibition in the OB strongly modulates frequency coupling of M/T cells.

## Results

### M/T cells differentially respond to different frequencies in odour stimuli

In order to probe the effect of temporal structure in odour stimuli on M/T cell activity, we recorded extracellular spiking activity using Neuronexus silicone probes (97 units, 6 mice) from the dorsal OB of anaesthetized mice while presenting 4 different odours (ethyl butyrate, 2-hexanone, amyl acetate and eucalyptol) at 5 different frequencies (2, 5, 10, 15 and 20Hz, Fig. 1A-B) using a high-speed odour delivery device we recently developed (Erskine et al., 2019). As M/T cells can respond to changes in air pressure due to mechanosensitivity of OSNs (Grosmaitre, Santarelli, Tan, Luo, & Ma, 2007), we offset changes in flow by presenting an odourless air stream from an additional valve following a temporal structure that operated anti-correlated to the odour valve. This resulted in approximately constant air flow profile throughout the odour presentation (Fig. 1B&C). The temporal odour delivery device (tODD) allowed for a reliable odour pulse presentation (Fig. 1C) with similar net volumes of odour for all the frequencies (P > 0.3 Welch’s t-test) (Fig. 1D). To control for responses to residual flow changes, we included ‘blank trials’, i.e. trials identical to the odour trials in temporal structure, except that both the valves were connected to a vial filled with mineral oil. Respiration was continuously monitored using a flow sensor placed in close proximity to the nostril contralateral to the recording hemisphere. To minimise sniff-cycle related variability (Shusterman et al., 2011) we triggered odour stimulation at the onset of inhalation.

**Figure 1:**
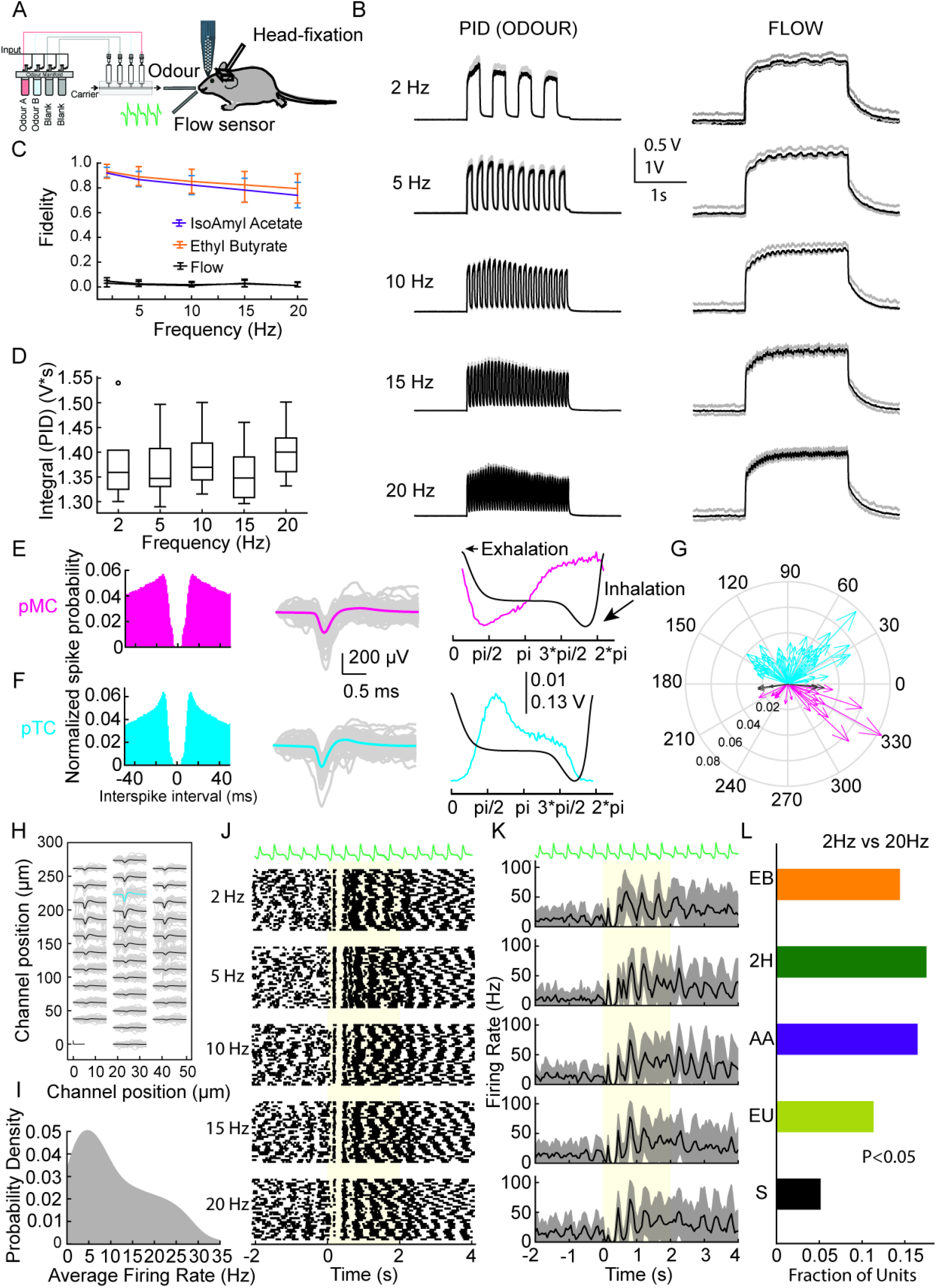
OB neurons encode odour frequency. **(A)** Schema of the unit recording experimental setup **(B)** Photo-Ionisation Detector (PID) signal (*left*) and airflow (*right*) measurements of Ethyl Butyrate stimuli at 5 different frequencies. **(C)** Fidelity of the odour stimuli and flow at different frequencies. **(D)** Integral values of the total odour for the 5 frequencies, shown in B, for 3 seconds after odour onset (2 seconds of odour and 1 second to allow full return to baseline). **(E)** Autocorrelogram from a ‘good’ example putative mitral cell (pMC) **(*left*)**, individual spike waveforms from the cell (grey) and its average waveform (magenta) **(*middle*)** and baseline spiking probability (magenta) overlaid on the respiration trace (black) **(*right*)**. Peak spiking probability coincides with inhalation phase of the animal. **(F)** Same as in (E) but for putative tufted cell (cyan) with peak firing probability coincides with exhalation of the animal. **(G)** Summary Phase plot displaying peak firing probability of all recorded units against phase of respiration (n=64; pTC (cyan), n=26; pMC (magenta) and n=7; unresolved (black)). **(H)** Average waveform for the pTC in (F) across all channels of recording probe. Average waveforms shown in black, with the cyan average indicating the channel with the largest average waveform. **(I)** Baseline firing rate distribution of all recorded units across all experiments. **(J)** Respiration trace **(top)** and raster plot for a single unit’s response to five odour frequencies. **(K)** Respiration trace **(top)** and average PSTH of the same unit’s responses in (J). **(L)** Fraction of units showing significant difference in firing rate in the first 500ms after odour onset between the 2Hz and 20Hz odour stimuli. The different colours represent the different odours used for all the units. (EB: Ethyl Butyrate; 2H: 2-Hexanone; AA: Isoamyl Acetate; EU: Eucalyptol; S: Shuffled)

A typical recording session yielded recordings from multiple clusters from a depth of 300-500 μm from the OB surface. The recorded clusters were classified either as ‘good’ (well isolated clusters), ‘MUA’ (multi-unit activity, clusters which contained spikes of physiological origin but from numerous cells), or ‘noise’ (clusters containing spikes of non-physiological origin, e.g. electrical interference, movement artefacts) based on their autocorrelograms (Fig. 1E-F *left*), waveforms (Fig. 1E-F *middle*), firing rate and amplitude stabilities. Corroborating previous findings (Ackels, Jordan, Schaefer, & Fukunaga, 2020; Fukunaga et al., 2012; Jordan, Fukunaga, Kollo, & Schaefer, 2018; Jordan, Kollo, & Schaefer, 2018) we observed that most of the units also coupled distinctly to either the inhalation (Fig. 1 E *right*, G) or exhalation phase (Fig. 1 F *right*, G) of the sniff cycle. From the entire population of recorded units, we estimated a total of 64 putative tufted cells (pTCs) and 26 putative mitral cells (pMCs) while 7 units could not be resolved into either of the class (Fig. 1G). The average baseline firing of the recorded units was found to be 11 ± 9 Hz (mean ± SD) (Fig 1I). Depending on the odour identity, between 13% and 50% of units displayed significant changes (>= 2 SD for 50ms from baseline) in their firing rates in response to the stimuli (Ethyl butyrate – 47/97; 2-Hexanone – 50/97; Amyl acetate – 43/97; Eucalyptol – 13/97) (Fig S1). Furthermore, a subset of units displayed visibly different spiking profiles in response to different frequency odour stimuli (Fig. 1J-K). Comparing e.g. activities of units between 2Hz and 20Hz stimuli we observed that a substantial number of units compared to the shuffled control showed significant difference in their activity in the first 500ms after odour onset. This was true for all the 4 odours tested (Ethyl butyrate – 14/97; 2-Hexanone – 18/97; Amyl acetate – 17/97; Eucalyptol – 12/97; Shuffled – 6/97, P<0.05 Unpaired t test) (Fig. 1L).

To examine the population level response to these different stimuli, we constructed response vectors by computing the cumulative spike count for all the 97 recorded units in the first 500ms post odour-onset and subtracting from it the spike counts of a blank trial. We trained linear classifiers on 80% of the data and tested on 20% to examine whether spiking activity obtained for different stimulus frequencies was linearly separable (Fig. 2A). We observed that when we used a 500ms rolling window with temporal steps of 10ms, a classifier could achieve accuracies (Ethyl butyrate: 55 ± 8%; 2-Hexanone: 51 ± 8%; Amyl acetate: 53 ± 8%; Eucalyptol: 45 ± 8) notably higher than what was obtained by training on shuffled data (20 ± 7%) (Fig. 2B). In addition, we trained classifiers on random subsets of unit responses, binned in a 500ms window from odour onset. The classifier accuracies increased with increasing number of units, with peak accuracies found for classifiers which had the full set of units available (Ethyl butyrate: 41 ± 5%; 2-Hexanone: 35 ± 5%; Amyl acetate: 46 ± 5%; Eucalyptol: 39 ± 5%) compared to shuffled data (20 ± 5%) (Fig. 2C). We then trained a series of classifiers to distinguish pairs selected from all possible combinations of odour frequencies and identities. We found that classifiers could readily distinguish responses for different stimulus frequencies well above chance (>0.5) for a given odour while performing even better while comparing responses for trials across different odours (Fig. 2D). Next, we withheld one trial of each type of stimuli and trained a classifier with all the remaining trials. We tested the classifier on the withheld trials to see both how well a classifier can distinguish trials, as well as explore the structure of the false classifications (Fig. 2E). For a given odour, the predictability of a frequency of stimulus reached well above chance (> 0.04). Furthermore, when comparing across the different odours, the predictability was almost perfect (Fig. 2E). Overall, this suggests that OB neurons can encode temporal structure present in odour stimuli at frequencies of at least up to 20Hz in their spiking pattern.

**Figure 2:**
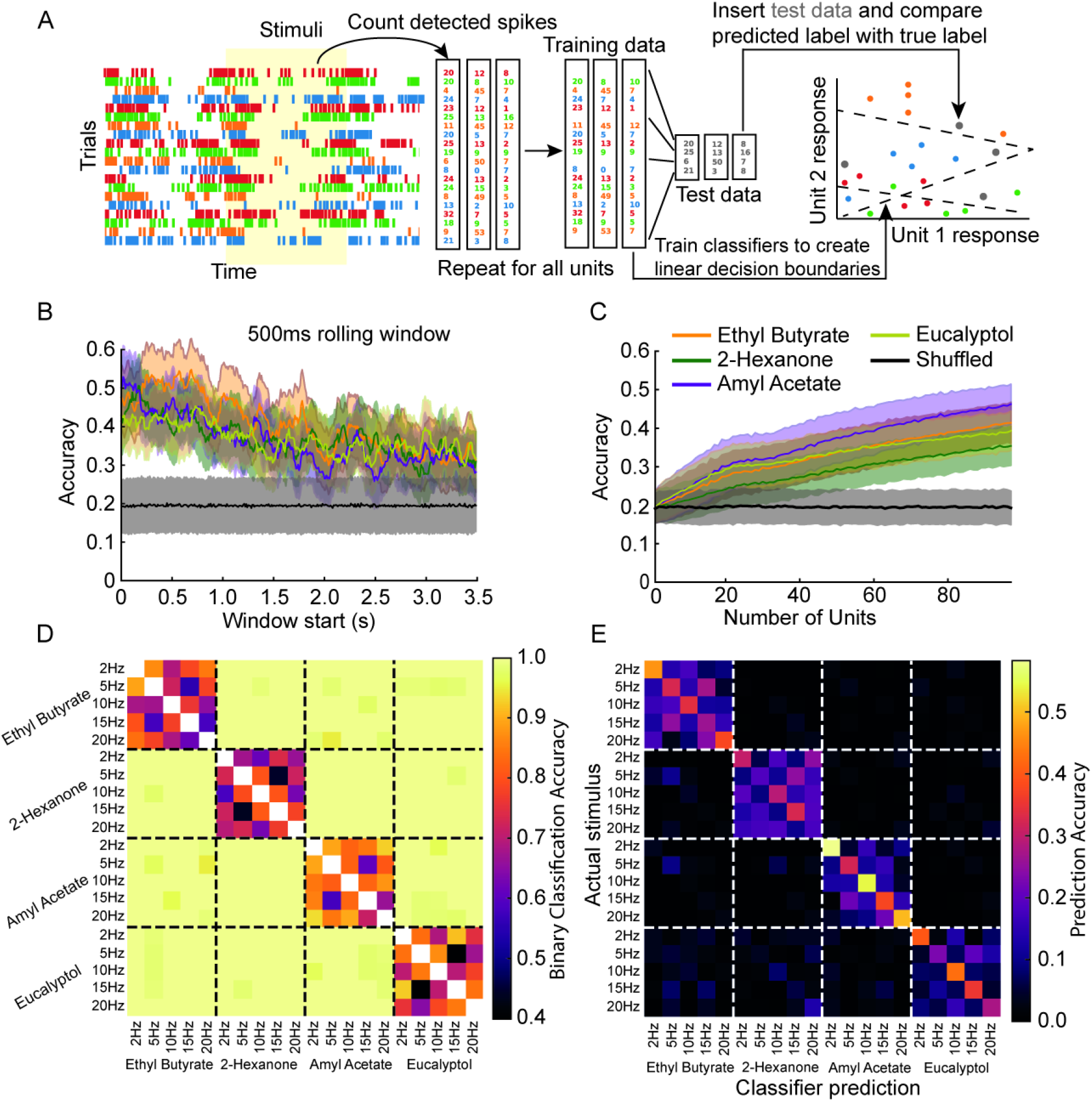
Classifiers to unit responses. **(A)** Schematic outlining of the procedure to train classifiers on unit spike times following an odour stimulus. **(B)** Average accuracy of linear SVM classifiers trained on a summed 500ms rolling window of detected unit spikes from different start times post odour onset. **(C)** Average accuracy of a set of linear SVM classifiers trained on the summed response from units 500ms from odour onset as the number of units available increases. **(D)** A matrix of all possible 1 vs. 1 comparisons between different trial types and the average accuracy of a linear SVM trained to distinguish between the two. Chance is 0.5 **(E)** A confusion matrix of linear fractional classifications for a set of classifiers trained to distinguish all trial types from one another. Y axis represents a trial’s true label and the x axis represents a classifier’s given label to a trial. Chance is 0.04.

### M/T cells follow odour stimulus both at 2 and 20Hz

To better understand the mechanism that gives rise to frequency dependent M/T cell spiking responses and to get insight into their subthreshold basis, we performed whole-cell recordings from M/T cells (Fig. 3A). To increase the probability of finding a responsive cell-odour pair and due to the time limitation and lower yield of reliable whole-cell recordings, we employed odour mixtures as stimuli (A: Ethyl butyrate + 2-Hexanone; B: Amyl acetate + Eucalyptol) and presented odours at only two frequencies, 2Hz and 20Hz (Fig. 3E). As with the unit recordings, stimuli were triggered at the onset of inhalation and blank trials were included. We recorded from 42 neurons in 25 mice at depths of 180-450 μm from the surface of the olfactory bulb (Fig. 3B). The neurons showed physiological RMP ranging from −38 mV to - 60 mV (Fig. 3C) and input resistance of 48-280 MΩ (Fig. 3D). These values are congruent with previous findings indicating that our recordings were largely from M/T cells (Fukunaga et al., 2012; Jordan, Fukunaga, et al., 2018; Jordan, Kollo, et al., 2018; Margrie, Brecht, & Sakmann, 2002; Margrie, Sakmann, & Urban, 2001; Margrie & Schaefer, 2003).

**Figure 3:**
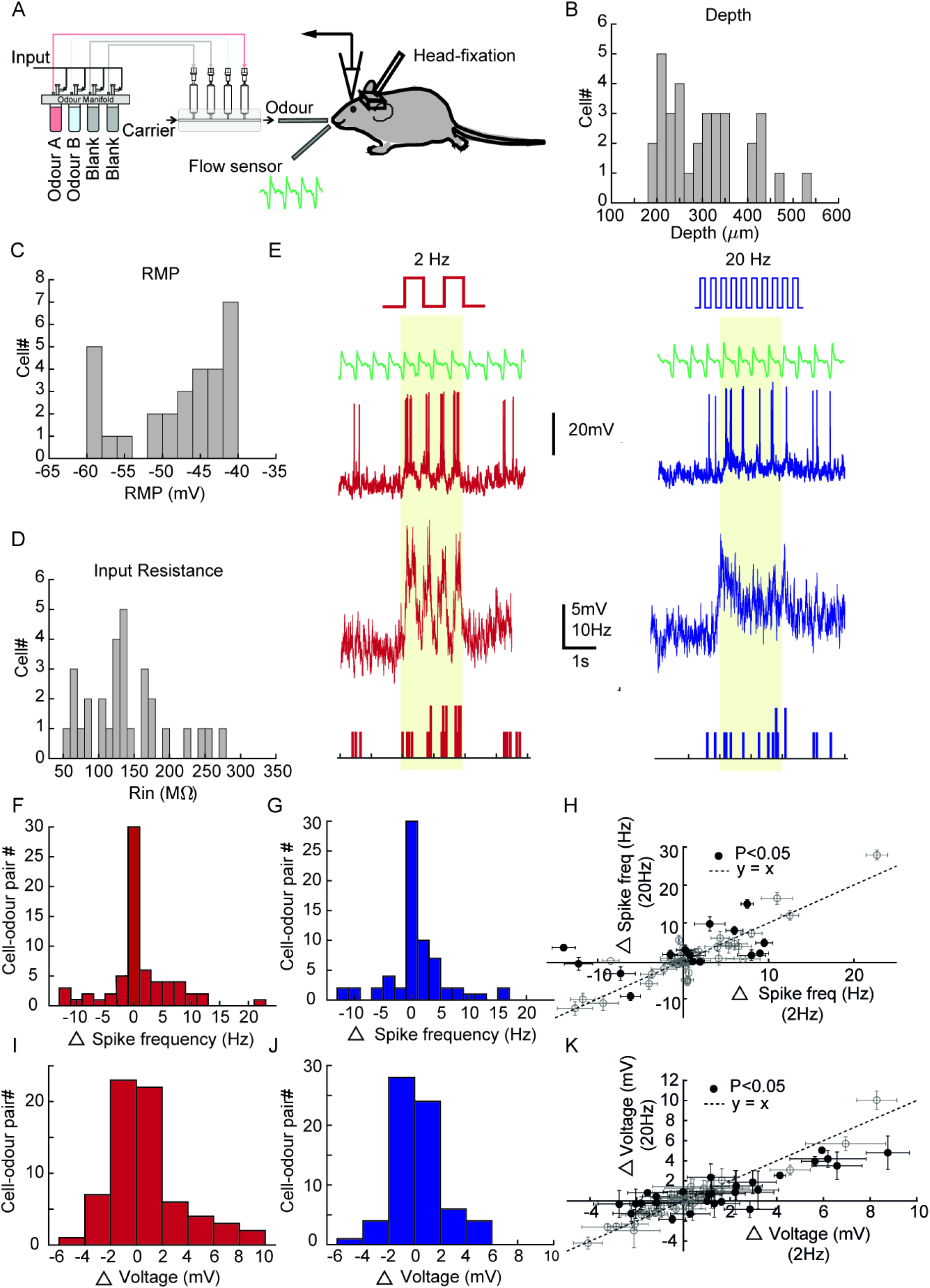
OB neurons respond to both 2 Hz and 20Hz temporally structured odour stimuli. **(A)** Schema of the whole-cell recording experimental setup. **(B)** Histogram representing the distribution of depth from the OB surface of all the recorded neurons, **(C)** resting membrane potential and **(D)** input resistance (n=42 cells, 25 mice) **(E)** Example recordings for 2Hz **(*left*)** and 20Hz **(*right*)** stimuli. From top to bottom, schema of odour stimulus, representative respiration trace, representative recording from a neuron, extracted V_m_ and PSTH. **(F)** Histogram of change in action potential frequency in the first 500ms compared to the baseline for 2Hz stimuli and **(G)** 20Hz stimuli. **(H)** Spike frequency change for 2Hz vs 20Hz for all cell-odour pairs. The hollow circles represent cells which showed statistically insignificant difference between the 2Hz and the 20Hz trials while the solid markers (15/68) represent cells showing significant difference (P<0.05, Two-tailed unpaired t-test). Each marker represents a cell and the error bars represent the SD obtained from all the trials. **(I)** Histogram of change in average V_m_ in the first 500ms compared to the baseline for 2Hz stimuli and **(J)** 20Hz stimuli. **(K)** Avg. Change in V_m_ for 2Hz vs. 20Hz. The markers and error bars have similar meaning as in (H) but for change in V_m_ from baseline. 27/68 cell odour pairs showed significant difference between the 2Hz and 20Hz trials (P<0.05, Two-tailed unpaired t-test).

Comparing average change in action potential firing frequency in the first 500ms after odour onset from the baseline between the 2Hz and 20Hz responses (3F-H), we observed that 15/68 cell-odour pairs behaved significantly different between the two cases (Fig. 3H, P<0.05, Two-tailed unpaired t-test). This is consistent with our findings from the unit recordings. To assess whether or how the subthreshold response of the recorded neurons for the 2Hz and 20Hz stimuli differ, we calculated the average change in membrane potential (Δ Voltage) during the first 500ms of the stimulus from the baseline (Fig. 3I-K). Overall, a larger number of cell-odour pairs (27/68) pairs showed significant differences between the responses to the two stimuli (Fig. 3K, P<0.05; Two-tailed unpaired t-test) compared to the suprathreshold response.

To quantify the coupling of membrane potential to the frequency of odour stimulation, we computed a coupling coefficient index (C*p*C, Fig. 4 A-F) for each individual cell (n = 42 cells, 70 cell-odour pairs, 25 mice, see Methods for details). In brief, to calculate C*p*C, the membrane potential was first baseline subtracted to minimise sniff-related membrane potential oscillations (Abraham et al., 2010). For a given cell-odour pair, C*p*C is then obtained by normalising the peak cross-correlation value for the odour response to that for the mineral-oil response (Fig 4 C-F), resulting in a C*p*C value > 0. A high C*p*C value indicates a cell that has a strong cell-odour frequency coupling. Further, a C*p*C > 1 indicates a cell’s response to the odour-frequency pair is stronger than that to a blank trial, and is not due to a response of a potential residual purely mechanical stimulus. A subset of the recorded neurons showed C*p*C >1 indicating substantial coupling to both 2Hz (35/70) (Fig. 4G) and 20Hz (25/70) (Fig. 4J). To assess statistical significance of this coupling measure, we shuffled the phases of the membrane potential before computing the shuffled control C*p*C (Fig. 4H&K).

**Figure 4:**
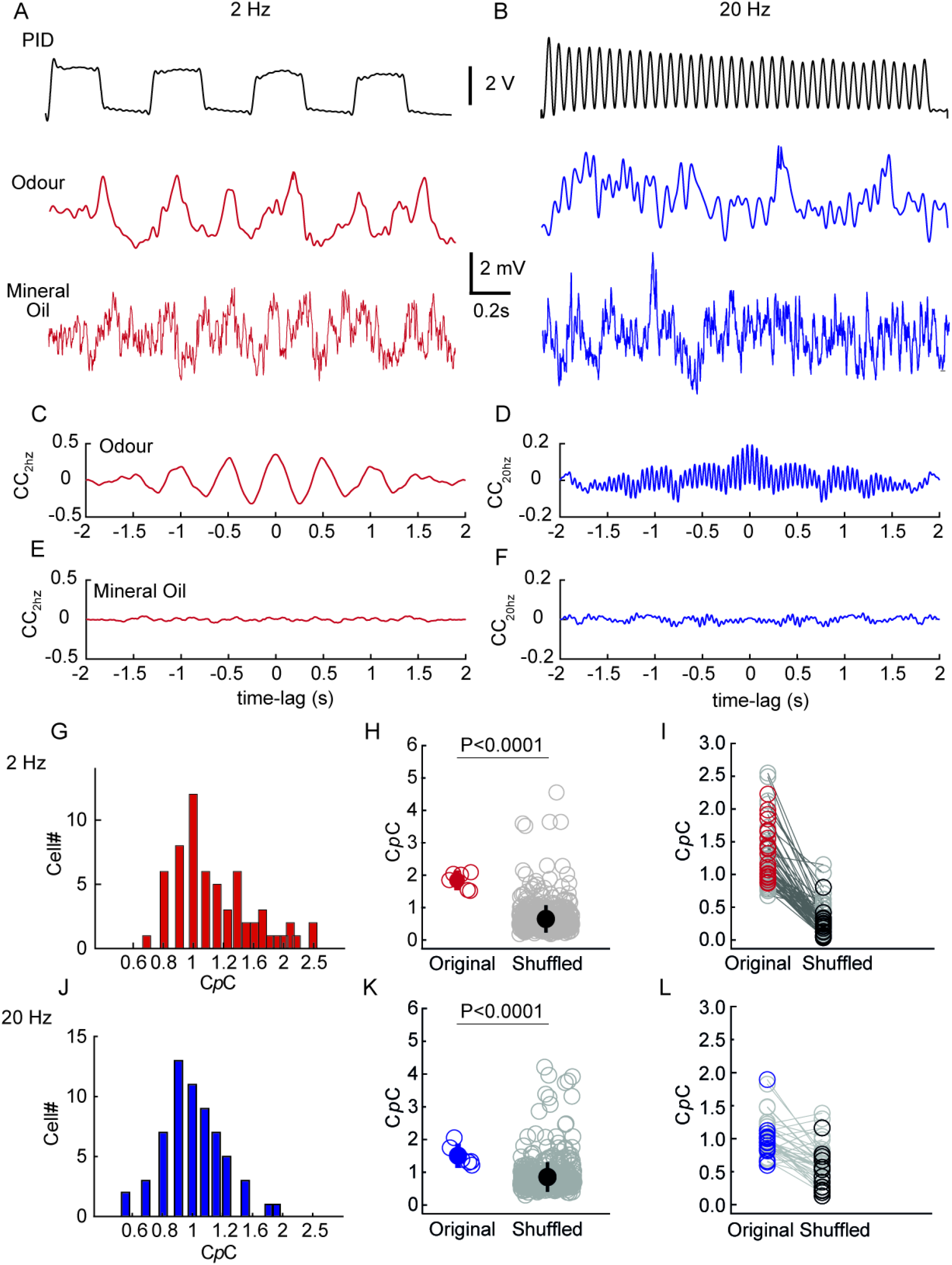
Olfactory bulb neurons can follow temporally structured odour stimuli. **(A) (*Top*)** A representative trace recorded using a PID of a 2Hz odour stimulus; **(*Middle*)** a representative V_m_ trace (action potentials clipped) recorded during 2Hz odour stimulus presentation; **(*Bottom*)** recording from the same neuron during 2Hz mineral oil stimulus presentation. **(B)** Same as in (A) but for a 20Hz stimulus from a different neuron. **(C)** Cross-correlation plot between 2Hz PID trace and 2Hz V_m_ odour trace; **(D)** 2Hz PID trace and 2Hz V_m_ mineral oil trace; **(E)** 20Hz PID trace and 20Hz V_m_ odour trace and **(F)** 20Hz PID trace and 20Hz V_m_ mineral oil trace. **(G)** Histogram of coupling co-efficient (C*p*C) for all cells for 2Hz responses (n=42 cells, 25 mice). Note a considerable fraction of neurons showing C*p*C > 1 indicating that they follow the temporal odour stimuli. **(H)** C*p*C estimated from the individual original trials (red hollow) and that from the shuffled control (grey) of 2Hz odour stimuli for a given cell. The filled circle represents the population average of C*p*C obtained from the original trials (red solid) and the shuffled controls (black). P<0.0001, Mann-Whitney test. **(I)** Original C*p*C vs shuffled C*p*C from all recorded cells. The grey markers (n=47/70) represent the cell-odour pair showing non-significant difference between the two while the coloured markers (23/70) represent the significantly different cells. Significance threshold was set at P<0.01 **(J)** Same as in G but for 20Hz responses **(K)** Same as in (H) but for 20Hz responses. (n=16/70, significant cell-odours) **(L)** Same as in (I) but for 20Hz responses.

Comparing the C*p*C between the original and the shuffled control, we observed that a substantial number of M/T cells indeed significantly coupled to both 2Hz (23/70 cell-odour pairs) (Fig. 4I) and 20Hz (16/70 cell-odour pairs) (Fig. 4L) (P<0.01, Two-tailed paired t-test). For a subset of recorded cells, we presented a third stimulus, a continuous odour (with no temporal structure) and estimated the C*p*C as before. As expected, C*p*Cs obtained from the 2Hz and 20Hz stimuli was significantly higher than that from the constant odour stimulus for a substantial portion of the recorded M/T cells (2Hz: 17/32 cell-odour pairs; 20 Hz: 13/26 cell-odour pairs; P<0.01, Two-tailed paired t-test, Fig. S2).

### Putative Tufted cells show higher C*p*C than putative Mitral cells

Spontaneous oscillation of membrane potential has been observed to be a reliable predictor of projection neuron type in the OB (Ackels et al., 2020; Fukunaga et al., 2012; Jordan, Fukunaga, et al., 2018). We classified the recorded neurons into 23 putative mitral cells (pMC) and 17 putative tufted cells (pTC) based on the phase locking of the spontaneous membrane potential to the respiration cycle of the mouse (Fig. 5A-B). Two of the cells could not be resolved. While overall similar, pTCs tended to couple more strongly to the odour stimuli than pMCs, reaching significance for the 20 Hz case (2 Hz: C*p*C_pTC_=1.32±0.48 (mean±SD), C*p*C_pMC_=1.24±0.41; P=0.64, KS test; 20 Hz: C*p*C_pTC_=1.12±0.29, C*p*C_pMC_=0.95±0.49; P = 0.04, KS test). This corroborated our depth-based classification (Fig. S4 E-F) where we observe that for the 20Hz cases, superficially located tufted cells showed higher C*p*C compared to the deeper mitral cells (Fig. S4F, P<0.05).

**Figure 5:**
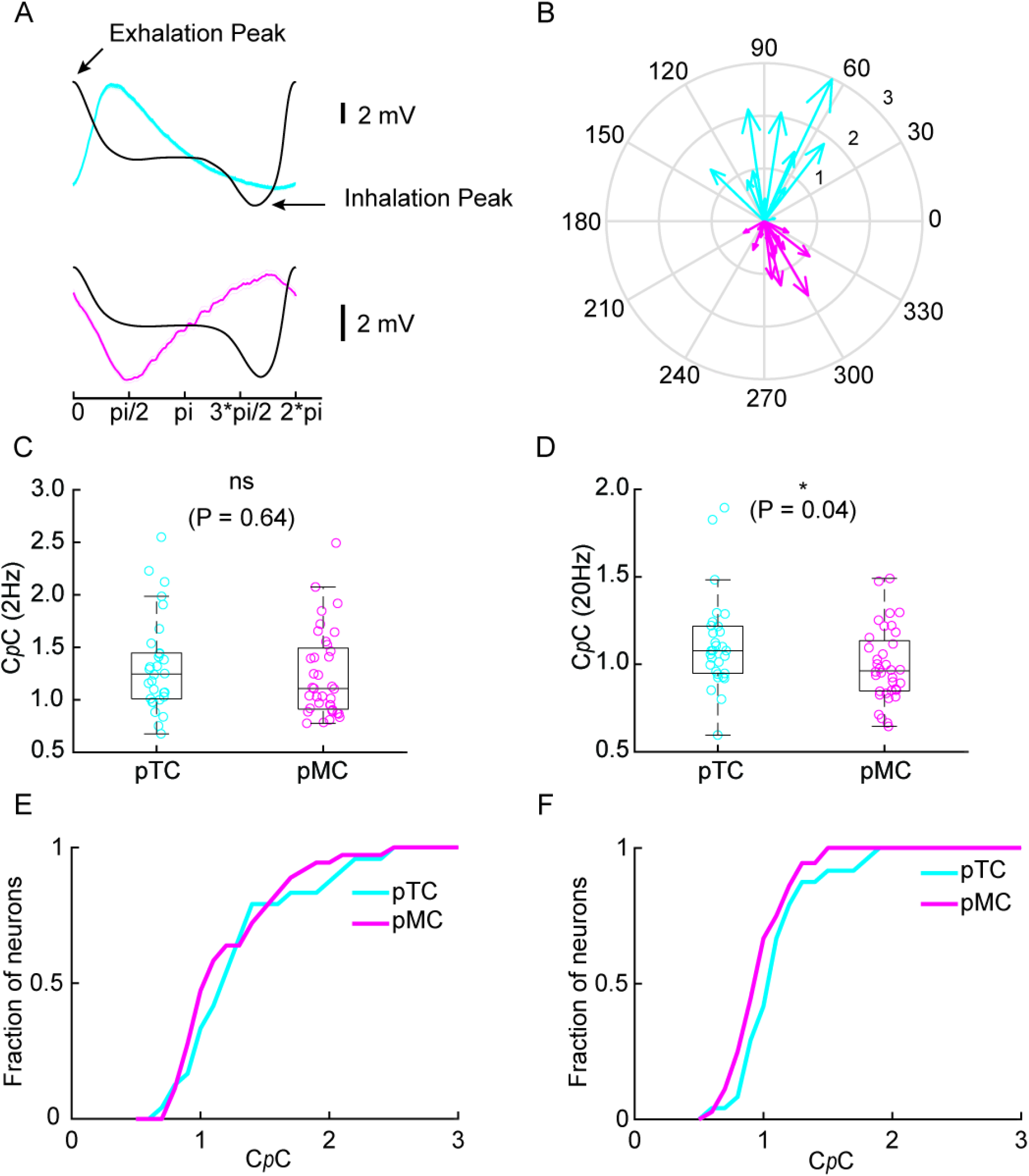
Putative Mitral and Tufted cells show similar probability of following both 2 and 20Hz stimulus. **(A) (*Top*)** Representative baseline average V_m_ trace from a putative tufted cell (cyan) overlaid with average sniff cycle (black). **(*Bottom*)** similar traces for a putative mitral cell (magenta). **(B)** Summary phase diagram of peak subthreshold oscillation phase plot (pTC; n=17, pMC; n=23). **(C)** C*p*C for all the recoded pTC and pMC for 2Hz showing no significant difference between the two population (P=0.64, KS test) while **(D)** pTCs show higher C*p*C than pMCs for 20Hz cases (P=0.04, KS test). **(E)** Cumulative histogram of C*p*C for all pTC and pMC for 2Hz and **(F)** 20Hz cases.

### C*p*C for odour mixtures can be linearly predicted from that of individual constituents

As described above, we presented two different odour stimuli at different frequencies (odour A and odour B). Comparing C*p*Cs for odour A with that displayed for odour B we noticed that these were tightly linked: M/T cells coupling well to odour A, also coupled well to odour B whereas M/T cells poorly coupling to one also coupled weakly to the other (Fig. 6A-B). This was the case for both 2Hz (Fig. 6C) and 20Hz (Fig. 6D) suggesting that the frequency coupling of a cell is independent of the odour presented. To further corroborate that C*p*C is an odour independent parameter, we probed M/T cells with a mixture of the two odours. If frequency-coupling is indeed odour-independent, a cell’s response to odour mixtures should be predictable from the C*p*C for the individual odour. To assess this, we first categorized recordings based on the direction of average change in membrane potential in the first 500ms after odour onset, and classified responses into 3 types: excitatory-inhibitory (Ex-In), excitatory-excitatory (Ex-Ex), and inhibitory-inhibitory (In-In) (Fig. 7A-B). Notably, we did not observe any significant difference between C*p*Cs for the different response types – neither for 2Hz (P = 0.54, One-way ANOVA; Fig. 7C) nor for 20Hz (P = 0.15, One-way ANOVA; Fig. 7D).

**Figure 6:**
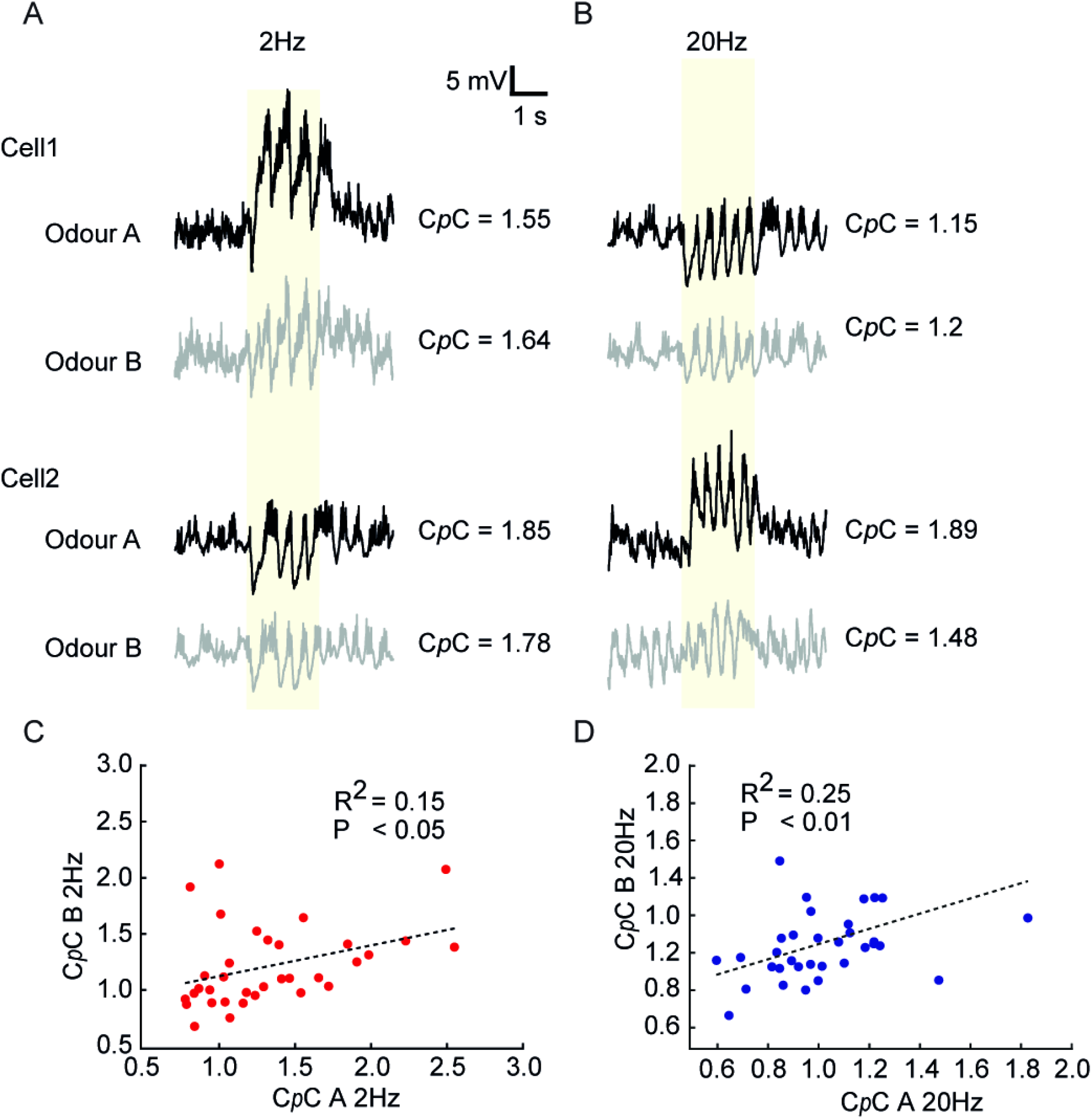
C*p*C is odour invariant. **(A)** Representative recordings from two cells responding to both odours A and B at 2 Hz. Note that C*p*C values are similar for the two odours. **(B)** As in (A) for 20Hz stimulation. **(C)** Plot of estimated C*p*C A vs C*p*C B from all the recorded cells for 2Hz and **(D)** 20Hz case. Note the strong correlation in both cases suggesting that coupling to frequency-modulated odour stimuli is odour-independent.

**Figure 7:**
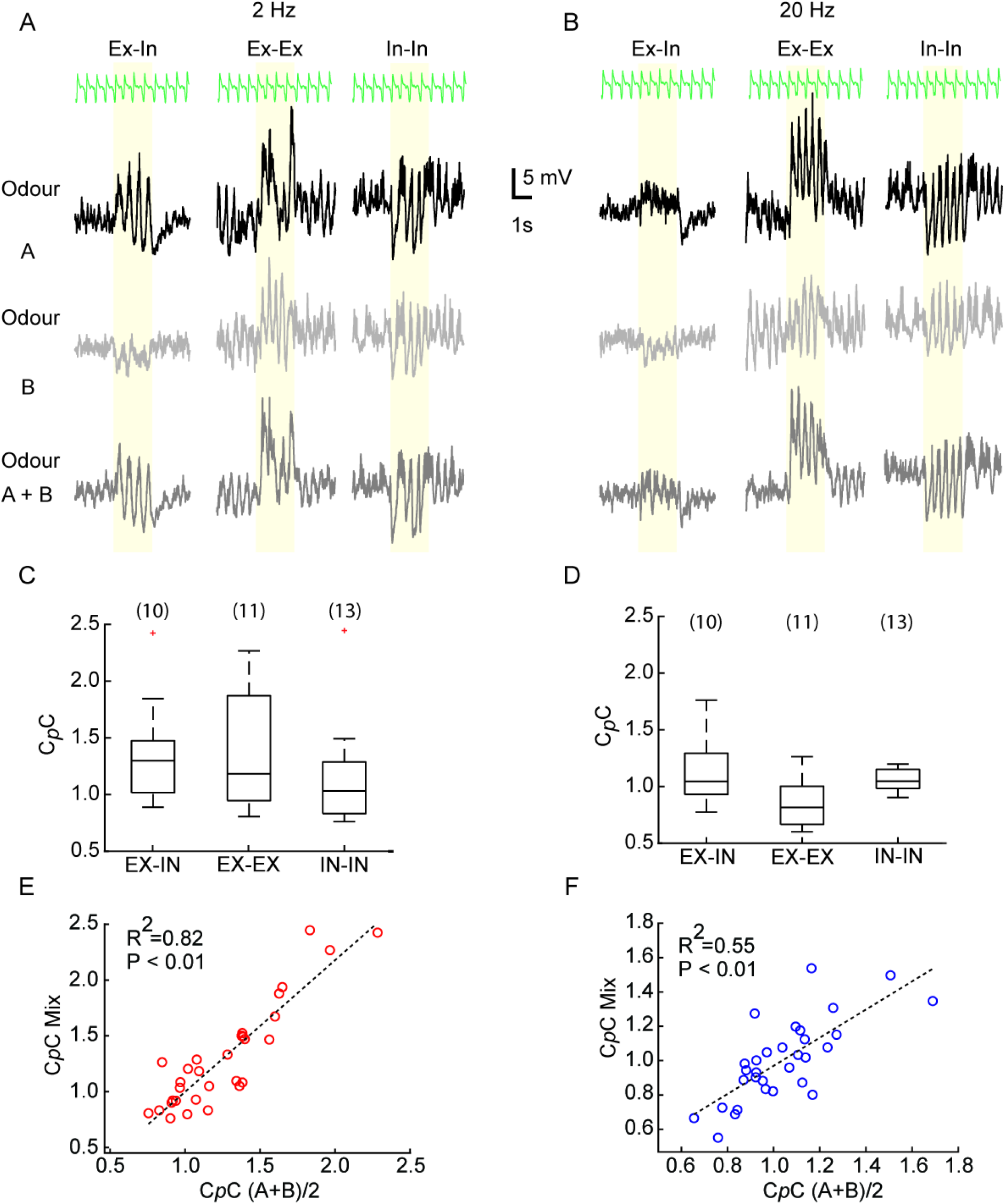
C*p*C for a mixture of odour can be linearly predicted from their individual component. **(A)** Representative recordings for 3 different types of mixture interaction; excitatory-inhibitory (Ex-In), excitatory-excitatory (Ex-Ex) and inhibitory-inhibitory (In-In). From *top* to *bottom* respiration trace, V_m_ for odour A, B and A+B mixture delivered at 2Hz. **(B)** as (A) but for 20Hz. **(C)** No significant difference in C*p*C between the 3 types of mixture responses for 2Hz and **(D)** 20Hz. The numbers in bracket are the number of cells. (1-way Annova, P=0.54 (2Hz); P=0.14 (20Hz)) **(E)** Computed C*p*C of odour (A+B)/2 vs actual C*p*C of mixture (A+B) for 2Hz cases and **(F)** 20Hz cases. Note linear regression can reliably predict the relation between calculated and estimated C*p*C (n=35, P<0.01).

Next, we tried to predict the C*p*C of a given cell to an odour mixture, based on the cell’s C*p*C obtained from the constituent odours. As predicted from the observation that C*p*C is largely cell-intrinsic and odour-independent, we observed that the C*p*C for a given cell-mixture pair could be reliably predicted from the cell-constituent pairs both for 2Hz (Fig. 7E) and 20Hz (Fig. 7F) (P<0.01). Notably, we observed that C*p*C was not correlated with the strength of odour response for a given cell-odour pair (Fig. S3).

### Influence of inhibition on C*p*C

Since our observations indicated that C*p*C is cell-intrinsic, independent of the odour presented (Fig. 6,7), we next asked whether a cell’s C*p*C was linked to intrinsic cellular or shaped by circuit properties. We observed that input resistance (Fig. S4A-B) and resting membrane potential (RMP, Fig. S4C-D) of a cell were not correlated with its C*p*C. C*p*C and depth showed different correlations for 2Hz and 20Hz. For the 2Hz cases we could not find any correlation between depth and C*p*C (Fig. S4E), while C*p*C decreased with depth for the 20Hz cases (Fig. S4F) (P<0.05). This suggests that tufted cells which are located more superficially than mitral cells might couple more strongly to high frequency (20 Hz) odour stimuli. This is consistent with our finding that pTCs showed significantly higher C*p*C to 20Hz than pMCs (Fig 5D).

To assess the role that circuit level inhibition may have on cellular C*p*C, we recorded from M/T cells as outlined previously, and then washed in a titrated mixture of 0.4mM gabazine and 2mM muscimol to cause “GABA_A_ clamping” (Fukunaga et al., 2012), blocking synaptic inhibition while providing sufficient unspecific, background inhibition to avoid epileptic discharge. Following a short period of change in membrane potential, the recorded neurons returned to approximately their original RMP (Fig. 8A). The input resistance (Fig. 8E) and RMP (Fig. 8H) did not change significantly in any of the neurons recorded. Under baseline conditions before GABA-clamping, the estimated C*p*C was significant compared to their shuffled controls for most of the recorded cell-odour pairs (13/15; 2Hz and 12/15; 20Hz, P<0.01, Two-tailed paired t-test). Post drug perfusion (with GABA_A_-clamp) all recorded cells were significantly coupled (15/15; 2 & 20Hz, P<0.01, Two-tailed paired t-test) (Fig. 8C-D). Furthermore, we noticed a significant change in C*p*C for most of the cell-odour pairs post drug treatment from their baseline values (n = 10/15; 2Hz and 9/15; 20Hz, P<0.01, Two-tailed paired t test) (Fig. 8F-G). Interestingly, this shift in C*p*C post GABA_A_ clamping happened in both directions with 5/15 cells showing a significant *decrease* and 5/15 a significant *increase* in their C*p*C values in the 2Hz case (Fig. 8F). For the 20Hz case, we observed that 2/15 cells show *decrease* while 7/15 show *increase* in C*p*C (Fig. 8G). Furthermore, we observed that cell-odour pairs which significantly showed an *increase* in C*p*C post GABA_A_ clamp from their baseline values, largely had initial C*p*C < 1 (2Hz: 0.947±0.076, 20Hz: 0.98±0.11 (mean±SD)), while for cell-odour pairs showing a decrease post GABA_A_ clamp had initial C*p*C > 1 (2Hz: 2.31±0.39, 20Hz: 1.3±0.07 (mean±SD)). Overall, our results indicate that the local inhibitory circuitry contributes significantly to determining a cell’s C*p*C.

**Figure 8:**
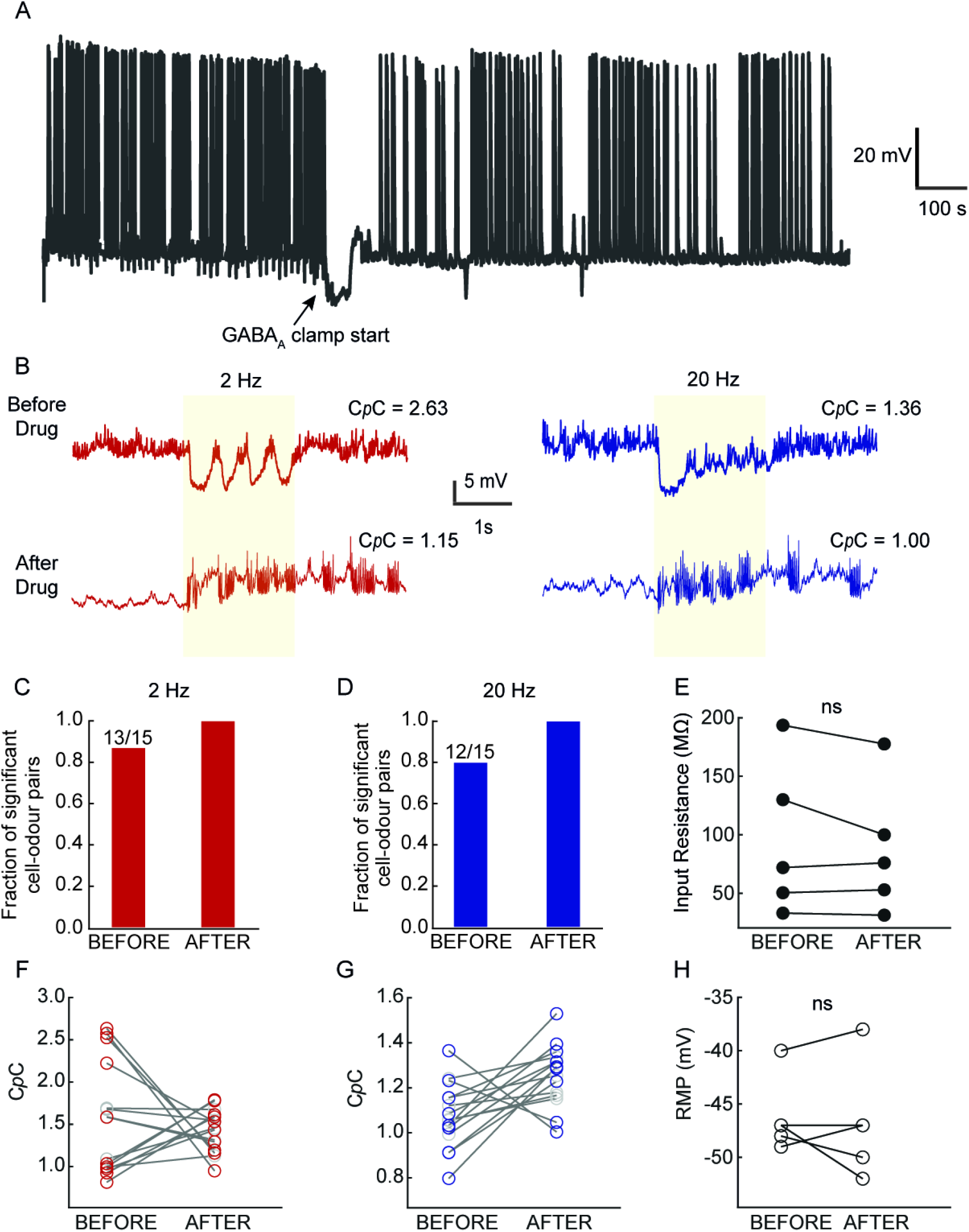
Influence of blocking inhibition on C*p*C. **(A)** Representative recording showing time of 2mM Muscimol + 0.4mM Gabazine infusion for GABA_A_ clamping. **(B)** Example V_m_ trace without drug **(*Top*)** and after 10 minutes from drug infusion point **(*Bottom*)** in a continuous recording for 2Hz (red) and 20Hz case (blue). **(C)** Fraction of cell-odour pairs showing significant C*p*C compared to their shuffled control before and after the drug infusion point for 2Hz stimuli. **(D)** Same as in (C) but for 20Hz cases. **(E)** Input resistance of the recorded neurons estimated both before and after drug infusion (n=5). No significant change was observed (Paired t test; P=0.272). **(F)** 10/15 cell-odour pairs (red) showed significant change in their C*p*C post GABA_A_ clamping while 5/15 (grey) showed no significant change in the 2Hz case (P<0.01, Two-tailed paired t-test). **(G)** 9/15 cell-odour pairs (blue) showed significant change in their C*p*C post GABA_A_ clamping while 6/15 (grey) showed no significant change in the 20Hz case (P<0.01, Two-tailed paired t-test). **(H)** RMP of the recorded cells did not change significantly due to GABA_A_ clamping

## Discussion

Olfaction research has largely used odour identity (chemical structure) and intensity as modulators of odour responses despite the presence of rich temporal structure in natural odour landscapes. Here we have shown that M/T cells in the OB *in vivo* can encode frequencies in odour stimuli as high as 20Hz. Furthermore, whole-cell recordings indicate that a subset of M/T cells significantly couple to frequencies of odour stimuli both at the sniff range and supra-sniff range in a largely odour-independent manner. We have demonstrated that odour frequency coupling capacity is similar between pMCs and pTCs populations at 2Hz, while at 20 Hz, pTCs couple more strongly than pMCs. Finally, while coupling capacity is independent of intrinsic properties, we observed that local inhibition strongly modulates a cell’s frequency coupling capacity. Overall, we show that the OB has the capacity to encode high frequency temporal patterns present in olfactory stimuli.

In mammals, respiration ensures a periodic sampling of olfactory stimuli which in turn is the main source of theta activity in the early olfactory areas (Macrides & Chorover, 1972; Margrie & Schaefer, 2003; Schaefer, Angelo, Spors, & Margrie, 2006). This causes rhythmic activity in M/T cells, even when devoid of odour stimuli (Connelly et al., 2015; Grosmaitre et al., 2007). Furthermore, the concentration of natural odour stimuli fluctuates in time (Celani et al., 2014; Erskine et al., 2019; Martinez & Moraud, 2013; Pannunzi & Nowotny, 2019; Riffell, Abrell, & Hildebrand, 2008) providing an extra layer of temporal information to the input signal of the OB. Altogether this creates temporally complex input signals for OSNs that are thought to carry information about the odour source (Celani et al., 2014; Erskine et al., 2019; Hopfield, 1991; Vergassola, Villermaux, & Shraiman, 2007). The fact that mice can discriminate optogenetically delivered excitation presented at different sniff phases (Smear et al., 2011) despite the relatively slow kinetics of signal transduction at the olfactory receptor neuron level (Ghatpande & Reisert, 2011) indicates the presence of circuit mechanisms (Gorur-Shandilya, Demir, Long, Clark, & Emonet, 2017) that might allow the early olfactory system to encode high frequency natural odour stimuli. Here we have shown that indeed the projection neurons in mouse OB can encode frequency in odour fluctuations and that local circuit inhibition plays an important role in this encoding.

### Functional implications

Naturally odours are carried by turbulent plumes of wind or water thus generating filamentous fluctuations of odour concentration (Celani et al., 2014) that can in principle contain information about the nature and location of odour sources (Erskine et al., 2019; Hopfield, 1991; Vergassola et al., 2007). Our observations here indicate that the M/T cells can encode information about odour fluctuations at frequencies from 2-20 Hz (Fig. 1–2, 4). Consistent with these results obtained from unit recordings, we observe that indeed a subset of neurons recorded intracellularly showed significantly different spiking and subthreshold membrane potential activity when presented with 2Hz or 20Hz fluctuating odour stimuli (Fig. 3H&K). Furthermore, our read-out parameter, C*p*C, provides an estimate of how strongly a given neuron can directly couple to a specific odour frequency. Both types of projection neurons showed overall similar frequency coupling to 2Hz (Fig. 5C) while tufted cells showed somewhat higher coupling than mitral cells to 20Hz (Fig. 5D and S4F). This might indicate that low frequency temporal information could be readily available to all the downstream cortical and sub-cortical areas while high frequency information cascades preferably to the anterior olfactory nucleus or anterior piriform cortex (Nagayama et al., 2010; Schneider & Scott, 1983; Scott, McBride, & Schneider, 1980). Further studies will be required to find exact structures in the downstream areas related to high frequency odour signal computation. The fact that blocking inhibition altered the frequency coupling capacity (Fig. 8) indicates that the inhibitory circuitry of the OB plays a strong role in encoding temporal features. This corroborates a previous finding in *Drosophila* projection neurons where blocking presynaptic inhibition altered response kinetics for temporally dynamic odour stimuli (Nagel et al., 2015). Additionally, we observe that post GABA_A_ clamping some of the cells displayed a decrease of C*p*C while other cells showed an increase (Fig. 8F-G). Further studies are required to pinpoint the precise inhibitory pathway which might be responsible for modulating frequency-coupling in M/T cells in a cell to cell basis and possibly identify subpopulations of MCs and TCs that encode specific temporal features.

Overall, here we provide one of the initial steps in understanding the capacity of the OB to encode temporally dynamic stimuli.

### Limitations of the study

All the recordings presented here were performed in anesthetized mice. Previous reports suggested that the behavioural state affects mitral cell firing properties (Kato, Chu, Isaacson, & Komiyama, 2012; Rinberg, Koulakov, & Gelperin, 2006). However, studies have suggested that M/T cell firing rates in awake conditions do not change but rather get redistributed over a breathing cycle (Gschwend, Beroud, & Carleton, 2012). Whole cell recordings from M/T cells indicate that membrane properties are largely similar between the two states (Kollo, Schmaltz, Abdelhamid, Fukunaga, & Schaefer, 2014), consistent with recent unit recording results (Bolding, Nagappan, Han, Wang, & Franks, 2020). It will nevertheless be important to repeat these experiments in awake mice to probe if the M/T cells hold to the same frequency coupling behaviour as that found in anaesthetised. Secondly, owing to the time limitation of reliable whole-cell recordings we could probe only two frequencies. Extracellular unit recording data partially alleviated this problem by investigating additional intermediate frequencies. Linear classifiers performed at accuracies well above chance level (Fig. 2), suggesting that M/T cells can encode several frequencies up to 20Hz. Therefore, it is likely that the frequency coupling capacity of subthreshold activity could span this range as well.

### *Temporally structured stimuli and pathophysiolog*y

In addition to better replicating naturalistic stimuli, temporally patterned sensory stimuli have been found to be advantageous in treating diseases. While direct electrical (and more recently optical) rhythmic deep brain stimulation is recognised as a possible treatment for a variety of neurodegenerative diseases (Benabid, Pollak, Louveau, Henry, & de Rougemont, 1987; Laxton et al., 2010; Zhang et al., 2019), it is only quite recent that temporally modulated sensory stimulation has been employed to in a similar manner. For example using 40 Hz visual and/or auditory stimuli has been found to help alleviate the amyloid burden from medial prefrontal cortex (Martorell et al., 2019). These reports strongly suggest that temporally structured sensory stimuli can be used as a tool to treat AD patients. A recent report indicating that humans can use temporal olfactory cues suggests this may be possible for olfaction as well (Perl, Nahum, Belelovsky, & Haddad, 2020). Therefore, temporally structured odour stimuli could offer an additional route for treatment for neurodegenerative disorders.

To conclude, in this study we report that M/T cells in the mouse olfactory bulb can encode temporal structure in odour stimuli and that their membrane potentials can follow frequencies of at least up to 20Hz. Furthermore, inhibition is key in generating this frequency-coupling.

## Material and Methods

### Animals

All animal procedures performed in this study were approved by the UK government (Home Office) and by Institutional Animal Welfare Ethical Review Panel. 5-8 weeks old C57/Bl6 males were used for the study.

### Reagents

All odours were obtained at highest purity available from Sigma-Aldrich, St. Louis MO, USA. Unless otherwise specified, odours were diluted 1:5 with mineral oil in 15 ml glass vials (27160-U, Sigma-Aldrich, St. Louis MO, USA).

### High-speed odour delivery device

A high speed odour delivery device was built as described previously (Erskine et al., 2019). Briefly, we connected 4 VHS valves (INKX0514750A, The Lee Company, Westbrook CT, USA) to odour containing 15ml glass vials (27160-U, Sigma-Aldrich, St. Louis MO, USA) through individual output filters (INMX0350000A, The Lee Company, Westbrook CT, USA). The vials were connected to a clean air supply (1L/min) through individual input flow controllers (AS1211F-M5-04, SMC, Tokyo, Japan). Each valve was controlled through a data acquisition module (National Instruments) controlled by a custom written script using Python software (PyPulse, PulseBoy; github.com/warnerwarner).

### In vivo electrophysiology

#### Surgical and experimental procedures

Prior to surgery all utilised surfaces and apparatus were sterilised with 1% trigene. 5-8 weeks old C57BL/6Jax mice were anaesthetised using a mixture of ketamine/xylazine (100mg/kg and 10mg/kg respectively) by injection inter-peritoneally (IP). Depth of anaesthesia was monitored throughout the procedure by testing the toe-pinch reflex. The fur over the skull and at the base of the neck was shaved away and the skin cleaned with 1% chlorhexidine scrub. Mice were then placed on a thermoregulator (DC Temperature Controller, FHC, ME USA) heat pad which was controlled using a feedback from a thermometer inserted rectally. While on the heat pad, the head of the animal was held in place with a set of ear bars. The scalp was incised and pulled away from the skull with 2 arterial clamps on each side of the incision. A custom head-fixation implant was attached to the base of the skull with medical super glue (Vetbond, 3M, Maplewood MN, USA) such that its most anterior point rested approximately 0.5 mm posterior to the bregma line. Dental cement (Paladur, Heraeus Kulzer GmbH, Hanau, Germany; Simplex Rapid Liquid, Associated Dental Products Ltd., Swindon, UK) was then applied around the edges of the implant to ensure firm adhesion to the skull. A craniotomy over the right olfactory bulb (approximately ~2mm diameter) was made with a dental drill (Success 40, Osada, Tokyo, Japan) and then immersed in ACSF (NaCl (125 mM), KCl (5 mM), HEPES (10 mM), pH adjusted to 7.4 with NaOH, MgSO_4_.7H_2_O (2 mM), CaCl_2_.2H_2_O (2 mM), glucose (10 mM)) before removing the skull with forceps. The dura was then peeled off using a bent 30G needle tip.

Following surgery, mice were transferred to a custom head-fixation apparatus with a heat-pad for recording. The animals were maintained at 37±0.5 °C.

#### Unit Recording

A Neuronexus poly3 probe was positioned above the OB craniotomy. An Ag/Ag^+^Cl^−^ reference coil was immersed in a well that was constructed of dental cement around the craniotomy. The reference wire was connected to both the ground and the reference of the amplifier board (RHD2132, Intan, CA, USA), which was connected (Omnetics, MN, USA) to a head-stage adapter (A32-OM32, Neuronexus, MI, USA). The probe, after zeroed at the OB surface, was advanced vertically into the dorsal OB at <4μm/s. This was continued until the deepest channels showed decrease in their recorded spikes indicating the end of the dorsal mitral cell layer. This was largely in the range of 400-600 μm from the brain surface. The signal from the probe was fed into a OpenEphys acquisition board (https://open-ephys.org/acquisition-system/eux9baf6a5s8tid06hk1mw5aafjdz1) and streamed through the accompanying GUI software (https://open-ephys.org/gui). The data was acquired at 30KHz and displayed both in a raw format, and a band pass filtered (300 - 6KHz) format. The band passed format was used primarily to visualise spikes across channels during the recording.

#### Odour stimulation (unit recordings)

Four odours (ethyl butyrate, 2-hexanone, isopentyl acetate, and eucalyptol) were diluted in mineral oil, as mentioned previous, in a ratio of 1:5.

Temporal structure of the odour stimulation was created using the VHS valves while blank valves helped maintain air flow to be constant through-out the stimulation period (Fig. 1B). The start of a stimulation was always triggered to the start of inhalation which was continuously monitored online using a flow sensor (AWM2000, Honeywell, NC, USA). A minimum of 8s inter-trial interval was given for all the experiments.

The onset pulse was passed to the OpenEphys acquisition board so that the trial trigger was recorded simultaneously with the neural data. 800 total trials were presented during the experiment, consisting of 32 repeats of 5 different frequencies for 4 odours and 1 blank. Each trial lasted 2 seconds and was spaced by a minimum of 8 seconds between the offset of one trial and the onset of the following trial.

#### Whole cell recording

Borosilicate pipettes (2×1.5 mm) were pulled and filled with (in mM) KMeSO_3_ (130), HEPES (10), KCl (7), ATP-Na_2_ (2), ATP-Mg (2), GTP-Na_x_ (0.5), EGTA (0.05) (pH = 7.3, osmolarity ~290 mOsm/kg). The OB surface was submerged with ACSF containing (in mM) NaCl (135), KCl (5.4), HEPES (5), MgCl_2_ (1), CaCl_2_ (1.8), (pH = 7.4 and ~300 mOsm/kg. Signals were amplified and low-pass filtered at 10 kHz using an Axoclamp 2B amplifier (Molecular Devices, USA) and digitized at 40 kHz using a Micro 1401 analogue to digital converter (Cambridge Electronic Design, UK).

After zeroing the pipette tip position at the OB surface, we advanced the tip to reach a depth of ~ 200 μm from the surface. Next, we stepped at 2μm/s to hunt for a cell in a similar manner as described before (Margrie et al., 2002; Margrie & Schaefer, 2003). Upon getting a successful hit, we released the positive pressure to achieve a gigaseal. The next gentle suction helped achieve the whole-cell configuration. We swiftly shifted to current-clamp mode to start a recording. Series resistance was compensated and monitored continuously during recording. Neurons showing series resistance >25 MΩ were discarded from further analysis.

The vertical depths of recorded neurons reported (e.g. Figure S4) are vertical distances from the brain surface. Respiration was recorded using a mass flow sensor (A3100, Honeywell, NC, USA) and digitized at 10KHz.

The GABA_A_ clamping experiments were performed as described before (Fukunaga et al., 2012). Briefly, muscimol and gabazine (Tocris Biosciences, Bristol, UK) were dissolved in ACSF to achieve a final concentration of 2mM (muscimol) and 0.4mM (gabazine). In a subset of experiments, this solution was superfused after ~10 minutes of recording under control conditions.

#### Odour stimulation (whole-cell recording)

Odours were presented as mixtures of monomolecular odorants mixed in 1:1 ratio which was eventually diluted in mineral oil in a 3:7 ratio. Odour A (ethyl butyrate + 2-hexanone) and B (isopentyl acetate + eucalyptol) were used for *in vivo* patch clamp experiments. Odour presentations were triggered on the onset of inhalation of the mouse as described for the unit recordings. The temporal structure of the odour pulses and the triggering of the blank valves were done as for the unit recording experiments described above. A minimum of 8s inter-trial interval was given for all the experiments.

### Analysis

#### Fidelity

Fidelity was defined here as the value of peak-to-trough of each square pulse normalised to the peak-to-baseline value. A fidelity of 1 therefore indicates that odour fully returns to baseline value between subsequent pulses, while a fidelity of ~ 0 for the flow indicates an almost continuous square pulse of air flow devoid of temporal structure.

#### Spike sorting

Kilosort2 (https://github.com/MouseLand/Kilosort2) was used to spikesort detected events into ‘clusters’. Clusters were then manually curated using phy2 (https://github.com/cortex-lab/phy) and assigned a ‘good’, ‘mua’, or ‘noise’ label depending on if they were considered to be made up of neural spikes (good and multi-unit activity), or false detections (noise). The clusters made up of spikes were further divided into good or mua dependent on if they are thought to be spikes from a well isolated single unit or not. A ‘good’ unit is characterised by a well-defined rest period in its auto-correlogram, a characteristic spike waveform, and a stable firing rate and spike amplitude (however these can both vary throughout the recording) (Fig. 1E-F). Only good clusters were used for further analysis.

#### Linear Classifier

97 clusters across 6 animals were grouped together and used for classification. Windows used to bin spikes varied both in size and in start relative to odour onset. Window starts span 0s to 3.99s from odour onset. The window sizes ranged from 10ms to 4s. All classifiers did not consider spikes from greater than 4s from odour onset. Therefore, a classifier with a 500ms window range could start between 0s and 3.5s, and a window range of 1s would use starts up to 3s from odour onset. This full range of starts and widths were used to determine both the time after odour onset that frequencies could be distinguished, and the time window required.

The data sets were always of dimensions 160 x 97 where 160 is the number of trials and 97 the number of clusters. The range of unit responses were scaled to have a mean response of 0 and a SD of 1. The data was split into a training (80%) and a test set (20%). The training set was used to train a linear Support Vector Machine (SVM) with a low regularisation parameter. The low regularisation parameter translates to less restrictions in weightings assigned to each cluster by the classifier. Once the classifiers had been trained, they were tested with the remaining 20% of trials. The trials to be saved for testing were picked at random. This training and testing were repeated 1000 times with a random selection of testing trials used each time. Classifiers were then trained on the same data but with their labels shuffled. To test how accuracy varied with number of clusters, random subsets of clusters were selected and used to train and test the classifiers.

Classifiers were then trained and tested on all 1v1 combinations of trials from the experimental data set. In this case a classifier was trained on all but two trials, one from each of the two trial types present in the training data. In this set chance was 50%. Finally, a series of classifiers were trained on all frequencies across all odours, with a single trial from every type being withheld for the test set. As there were 20 total trial types (5 frequencies with 4 odours) chance was 4%.

#### Change in membrane potential

The raw recordings were spike-clipped using custom script in spike2 (Cambridge Electronic Design, UK). They are then stored into MATLAB (Mathworks, USA) readable files for further analysis.

All the recordings used have been baseline subtracted to rule out the effect of sniff related background membrane potential oscillations. This was done as described previously (Abraham et al., 2010). Briefly, stretches of baseline period were collated after matching the sniff-phase to that during the actual odour presentation. The membrane potential associated with these baseline periods was averaged to make a generic baseline trace for every cell. This was then subtracted from all the recorded traces during the odour-stimulation period to create a baseline subtracted trace.

For calculating the average change in membrane potential for 2 and 20Hz; we averaged the membrane potential in a 2s period pre odour onset (Vm_base_). Next, we averaged the membrane potential in the first 500ms (~ 2 sniffs) post odour onset (Vmod_our500_) and subtracted from the baseline average voltage. In short;

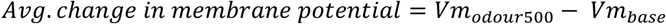

#### Change in spike frequency

Action potentials are counted in the raw data and converted into spike frequency in bins of 50ms. Bar plot of the spike frequency yields PSTH plots in Fig. 3E. Further, we calculate the average spike frequency in 2s before onset (FR_base_) and 500ms post onset (FR_odour500_) and eventually subtracting from each other to calculate the net change in spike frequency. In short;

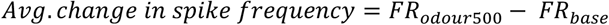

#### Frequency-coupling coefficient estimation

Baseline subtracted membrane potential traces for every odour and frequency were collected (Fig. 4A *middle*). PID traces recorded for the 2Hz and 20Hz odour stimulation was also averaged over 10 different trials (Fig. 4A *top*). Next, we cross-correlated the PID trace and all the individual baseline subtracted traces. This was repeated for all the trials for a given odour and frequency. We selected the peak correlation (CCodour_2Hz_ or CCodour_20Hz_) for all the trials. Similarly, we repeated the same exercise for the control blank stimulus which was also delivered at 2 and 20Hz and obtained a CCblank_2Hz_ or CCblank_20Hz_. Next, we normalized the CCodour with the CCblank for the respective frequencies and averaged them over all the trials to achieve a frequency-coupling coefficient (C*p*C) for a given cell-odour pair. In short for a given cell-odour pair:

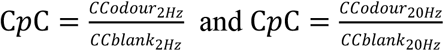

#### Shuffled control

For every recorded trial a set of 100 shuffled controls were created by shuffling the phase of the signal while maintaining the power similar to that of the original trace. We used custom written script in MATLAB for this process. A Fast-Fourier transform was done on the baseline-subtracted trace followed by separating the amplitude and phase of the signals. Next, the 0 Hz components were isolated while the other components were used for shuffling. Next, the phase and amplitude pairs for one half of the signal was shuffled in a pseudo random method using the ‘randperm’ function in MATLAB (Mathworks, USA). Next, conjugates of the thus shuffled combinations were created and concatenated with the previous shuffled half. This created a set of random combination of amplitude and phase pair of the same number of sampling points. Next, the signal was recreated using thus obtained shuffled amplitude and phase pairs and the 0Hz components. This was done 100 times for every cell-odour pair and frequency. The C*p*C was then estimated from each of these individual shuffled control in the same manner as described above.

## Acknowledgements

We thank the animal facilities at National Institute for Medical Research and the Francis Crick Institute for animal care and technical assistance. We thank the mechanical and electronic workshop (A. Ling, A. Hurst, M. Stopps); and S. Ghatak and members of the Sensory circuits and Neurotechnology laboratory for comments on earlier versions of the manuscript. This work was supported by the Francis Crick Institute which receives its core funding from Cancer Research UK (FC001153), the UK Medical Research Council (FC001153), and the Wellcome Trust (FC001153); by the UK Medical Research Council (grant reference MC_UP_1202/5); a Wellcome Trust Investigator grant to ATS (110174/Z/15/Z).

## Author Contributions

ATS and DD conceived the project, DD, ATS and TW designed experiments with inputs from AE; DD and TW performed experiments and analysed data, AE contributed tools and to experimental design; DD and ATS wrote the manuscript with input from all authors.

## Author Information

The authors declare no competing financial interests. Readers are welcome to comment on the online version of the paper. Correspondence and requests for materials should be addressed to DD and ATS.

